# The Loci of Insect Phenotypic Evolution

**DOI:** 10.1101/2022.11.29.518325

**Authors:** Virginie Courtier-Orgogozo

**Affiliations:** Université Paris Cité, CNRS, Institut Jacques Monod, 75013 Paris, France

**Keywords:** Genetics, Gephebase, compilation, natural evolution, insects, hotspot gene, repeated evolution, polygenic, recombination, standing variation

## Abstract

Insects are important elements of terrestrial ecosystems because they pollinate plants, destroy crops, transmit diseases to livestock and humans, and are important components of food chains. Here I used Gephebase, a manually curated database of genetic variants associated with natural and domesticated trait variation, to explore current knowledge about the genes and the mutations known to contribute to natural phenotypic variation in insects. Analysis of over 600 mutations reveals that data are concentrated towards certain species and traits and that experimental approaches have changed over time. The distribution of coding and cis-regulatory changes varies with traits, experimental approaches and identified gene loci. Recent studies highlight the important role of standing variation, repeated mutations in hotspot genes, recombination, inversions, and introgression.

**Highlights:**

- Gephebase compiles more than 600 genes and mutations contributing to insect natural variation
- Our genetic knowledge is biased towards certain traits and insect species
- Experimental approaches and studied insect species have changed over the years
- The relative distribution of coding and cis-regulatory mutations varies with traits and genes
- Clusters of causal mutations are more frequently found in insects than in other organisms

## Introduction

Insects impact humans as pollinators, sources of diverse products (honey, silk, etc.), sources of wonder (butterfly wings), crop pests, and vectors of pathogens for humans and domesticated plants and animals. Characterizing the genetic basis of insect phenotypic diversity is important not only for fundamental research to refine our understanding of evolution but also to better apply knowledge relevant to ecology, agriculture, and public health. For example, the identification of the genes underlying natural insect phenotypic variation can help to uncover promising targets and molecular pathways for disease control, pest control, and bioconservation.

Genetic studies of insect evolution began more than 100 years ago, with the pioneering work of Alfred H. Sturtevant on the sister species *Drosophila melanogaster* and *D. simulans* in Thomas H. Morgan’s laboratory [1]. Since then, the model organism *D. melanogaster* has been at the forefront of genetic discoveries, due to its ease of manipulation, the existence of powerful experimental tools, and accumulated knowledge from the work of thousands of *Drosophila* researchers across the world [2,3]. In the last thirty years, the development of increasingly efficient sequencing methods together with CRISPR/Cas-9 gene editing technology has broadened the range of insect species available for genetic studies.

More than 1000 insect genomes are now registered at the National Center for Biotechnology Information [4]. Most insect genomes and gene data are stored in integrated databases, such as i5k Workspace@NAL [5] and InsectBase [6], or in species-specific databases [4]. These databases are genome-centered and they do not provide details about the mutations and the genes that have been associated with natural phenotypic evolution. Gephebase (https://www.gephebase.org/) is a unique database that encompasses all Eukaryotes (mostly animals, yeasts, and plants) and consolidates published data about the genes and the mutations contributing to evolutionary changes [7]. Here, we used this database to review our current knowledge of the genetic basis of natural phenotypic evolution in insects.

Gephebase is manually curated and aims to document the genetic changes that have contributed to natural evolutionary processes. It does not include human clinical traits or mutations generated by random or directed mutagenesis. This compilation was initiated in 2008 as an Excel sheet synthesizing 331 entries [8] and it now contains more than 2500 entries. Each entry in Gephebase corresponds to a pair of alleles (from two different species or from two individuals of the same species) associated with phenotypic variation, either observed in nature or following domestication or experimental evolution in the laboratory. Each entry includes information about the species or population, the type of trait, the gene, the nature of the mutation(s), whether they represent null alleles, and the articles that reported it.

We gathered 600 insect entries in Gephebase and here we examine the current status of our knowledge about the genes and the mutations contributing to natural phenotypic variation in insects.

## Materials and Methods

### Literature Curation in Gephebase

Data is manually curated in Gephebase by a team of fewer than a dozen researchers. Newly published studies are found by this team using Pubmed and Google Scholar searches, references present in curated articles and reviews [9–14], and suggestions from colleagues that are collected via the “Suggest an Article” button on https://www.gephebase.org/. Gephebase compiles studies where the genotype–phenotype association is well supported [7]. For example, genetic loci are included only if there is functional evidence supporting the effect of the mutation on the phenotype of interest. One entry corresponds to a single mutation or a group of linked mutations within a single gene, where each mutation has been associated with a phenotypic effect. When the same mutation has occurred several times independently within the same species, distinct Gephebase entries are created.

Gephebase is up-to-date for all Eukaryotes for studies published prior to 2013. Because of an explosion of relevant studies in recent years, after 2013, we focused mostly on color variation in vertebrates (Gephebase “Trait” Advanced Search field = “Coloration”) [15] and on phenotypic variation in insects (Gephebase “Taxon and synonyms” Advanced Search field = “Insecta”). Compared to our last published report from 2020 [7], Gephebase contains more than 200 additional insect entries. The download of Gephebase data on 27 November 2022 (Supplementary File 1), which was used for the present study, can be considered as a comprehensive compilation of the genes and mutations contributing to insect natural phenotypic evolution that have been reported in the literature up to October 2022.

### Meta-analysis and figure preparation

Statistics and figures were produced using RStudio 2022.07.2+576 and R version 4.2.1 (2022-06-23).

## Results and Discussion

### Our genetic knowledge is biased towards certain species and traits

As of November 2022, Gephebase contains 694 mutations reported in the literature that contribute to phenotypic evolution in insects (Fig. 1). These mutations are associated with 41 traits and the two most represented traits, Xenobiotic resistance and Coloration, correspond to 61% and 16% of the total number of mutations, respectively (Fig. 2A). Very few traits in immature stages (embryo, larva, nymph) have been associated with mutations in specific genes. Most of the mutations present in Gephebase affect adult characters. Among the 108 mutations associated with Coloration, most (89%) relate to adult stages and only 6% and 4% to nymphal and larval coloration, respectively, and none to embryos. Only 11 mutations underlying behavioral variation (1.6% of all mutations reported in insects) have been identified (Fig. 2C). In general, behavior is more difficult to study and more variable than morphology and physiology, which probably accounts for the small number of studies identifying genetic causes of behavioral variation.

**Fig. 1.**
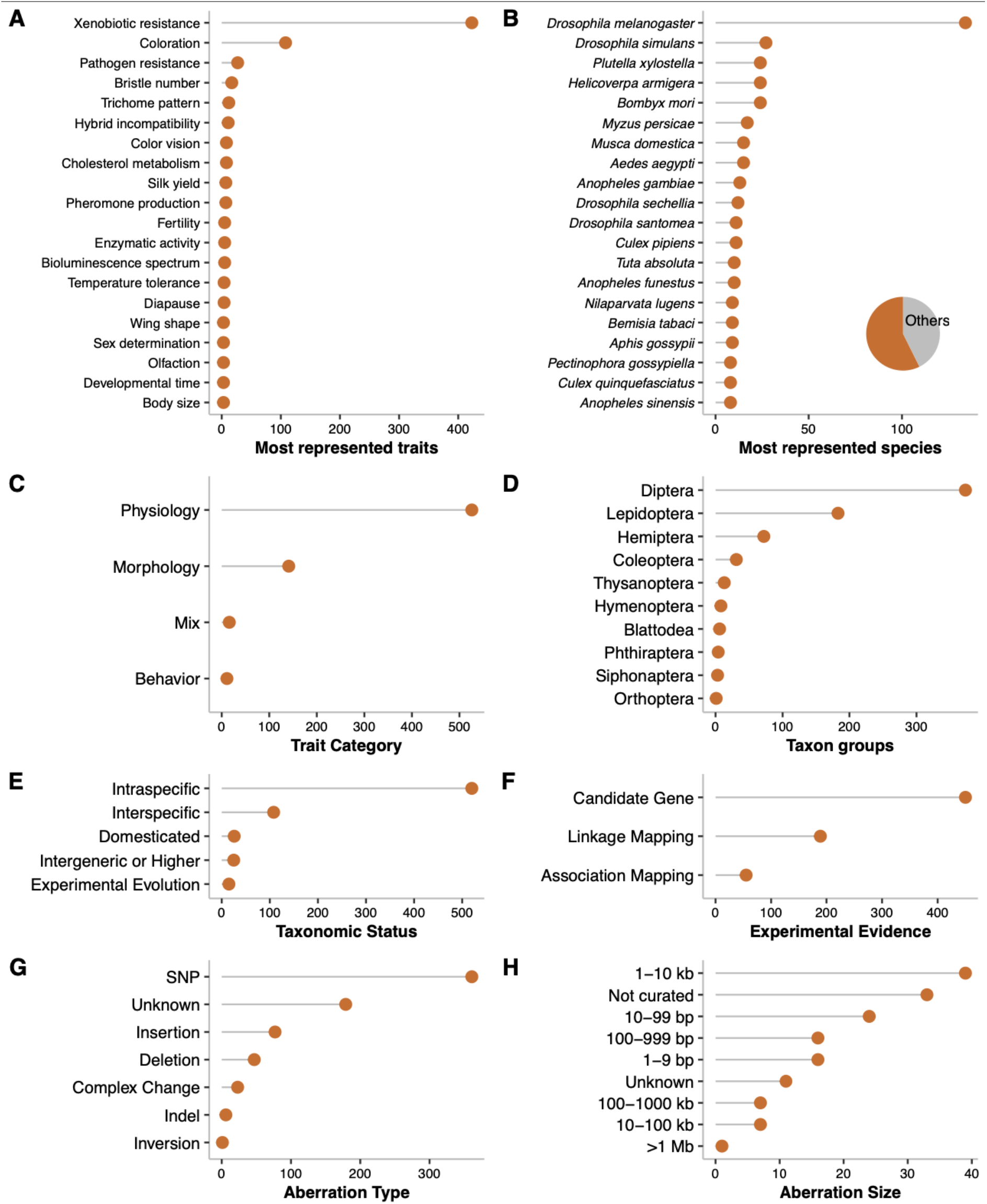
Distribution of the 694 mutations present in Gephebase that underlie natural phenotypic variation in insects. The x-axis indicates the total number of mutations for each category. (A) Twenty most represented traits. (B) Twenty most represented species. (C) Trait categories. (D) Taxonomic groups. (E) Taxonomic status. (F) Type of empirical evidence. (G) Aberration type. Indels are cases involving either a deletion or an insertion and where the direction of change is unclear. (H) Aberration size. Mutations classified as ‘SNP’ or ‘Unknown’ in (G) are not represented in H. The ‘SNP’ (single nucleotide polymorphism) category includes nucleotide substitutions between species for interspecific and intergeneric changes.

**Fig. 2.**
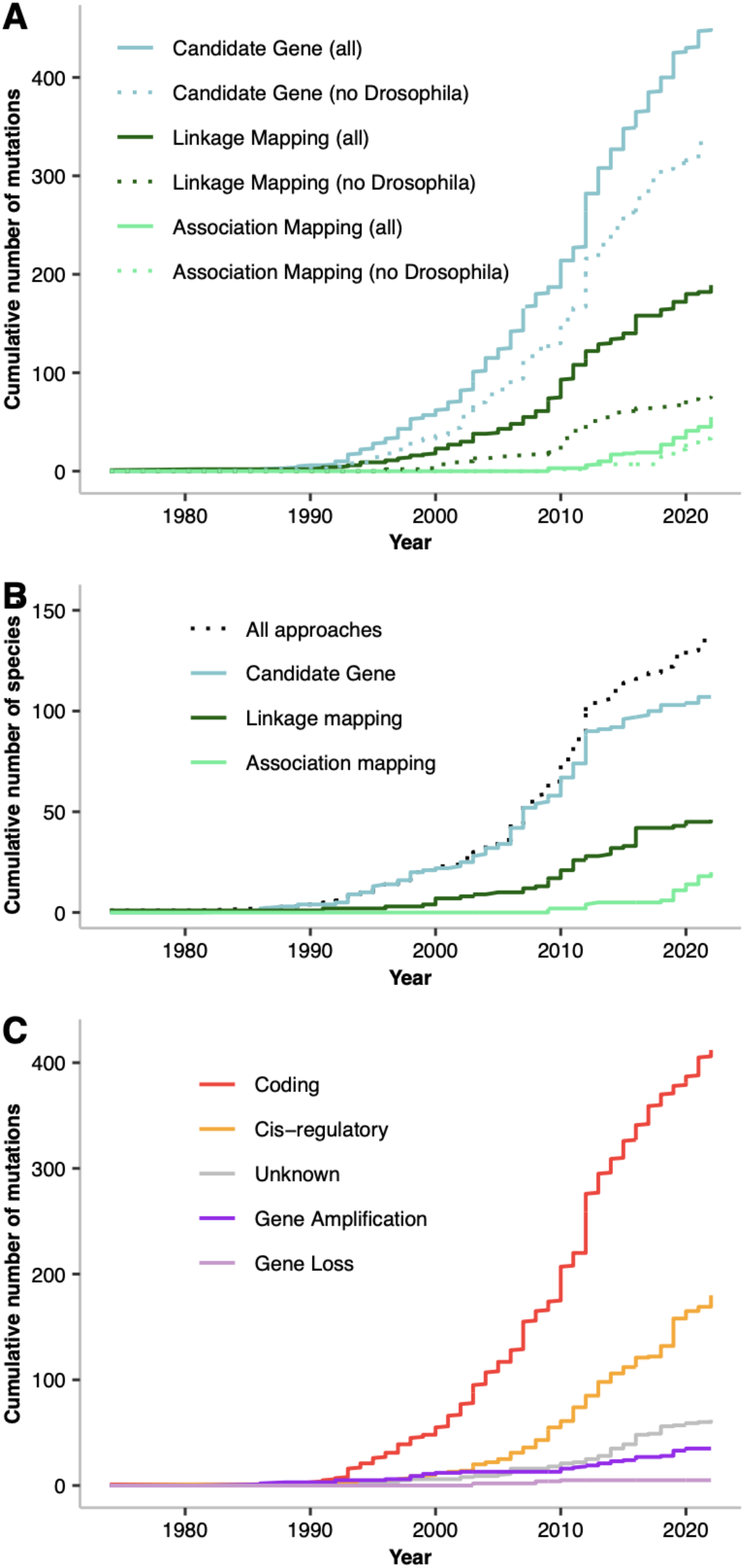
Cumulative number of mutations that have been identified over the years as responsible for insect phenotypic evolution and cumulative number over time of species for which genes and mutations have been shown to contribute to insect phenotypic evolution. (A) Cumulative number of mutations for the three Experimental approaches, when considering all species, or non Drosophila species. (B) Cumulative number of species for the three experimental approaches or for all approaches (dotted line). (C) Cumulative number of mutations for each molecular type: coding, cis-regulatory, gene amplification, gene loss and unknown (i.e. the gene locus was identified but not the molecular type of the mutation(s)).

Evolutionary relevant mutations have been characterized in a total of 158 insect species. The top species with most identified mutations (19%) is *Drosophila melanogaster*, as expected, followed by its sister species *D. simulans*, the cotton bollworm *Helicoverpa armigera* and the domestic silk moth *Bombyx mori* (Fig. 2B). Current data is biased towards Diptera (54%) and Lepidoptera (26%) (Fig. 2D). Although Coleoptera is the most speciose insect order, including about 38% of all known insect species [16], it contributes to only 4% of the mutations compiled in Gephebase for insects.

Gephebase mostly comprises intraspecific changes (75% of the mutations) (Fig. 2E). Indeed, most of the available experimental approaches to identify and then validate the genetic loci underlying phenotypic variation are dedicated to the comparison of closely related species or individuals from the same species that differ in particular traits.

Overall, our genetic knowledge is biased towards adult traits, xenobiotic resistance, coloration patterns, intraspecific changes, and a small set of species. It is not representative of the huge diversity of forms and life styles across the insect world.

### Genetic approaches and study species change over time

Genetic loci of evolution are typically identified via three approaches. The most highly represented one for insects is the candidate gene approach (Fig. 2A). The linkage mapping approach (QTL mapping) started to yield genes and mutations contributing to insect evolution in the 1990s for *Drosophila* flies and about ten years later for other insects. Then, around 2010, association mapping studies started to appear in the literature with successful identification of genes underlying insect evolution. Association mapping allows the study of natural insect populations when crosses in the laboratory are not possible. As of today, the mutations identified by association mapping are in Lepidoptera (33/55 mutations), *D. melanogaster* (12/55), *Anopheles* mosquitoes (4/55), the lady beetle *Harmonia axyridis* (3/55), the aphid *Hormaphis cornu* (1/55) and the bees *Bombus melanopygus* and *Lasioglossum albipes* (2/55). Gephebase contains more mutations identified via linkage mapping than via association mapping (Fig. 2A). The number of species for which trait variation has been mapped to precise genes keeps increasing and is currently higher for linkage mapping than for association mapping (Fig. 2B). However, with the decreasing costs of sequencing, I believe that association studies will become more prevalent in future discoveries of mutations contributing to insect evolution.

Identification of the genes underlying insect variation have been boosted in recent years by the advent of CRISPR/Cas9 (Clustered Regularly Interspaced Short Palindromic Repeats) biotechnology, which allows introduction of mutations of interest in practically any insect genome, provided that embryos or gonads can be injected with a CRISPR/Cas9 mix and resulting individuals obtained [17]. In the last five years, CRISPR/Cas9 has thus been used to validate in *D. melanogaster* the effect of more than 20 nucleotides changes associated with insecticide resistance in various insect species [18]. Furthermore, for a substitution identified by genome-wide association in the cotton bollworm *Helicoverpa armigera*, the mutation was introduced in a susceptible strain of the pest species and conferred a 125-fold resistance, confirming the effect of the identified substitution [19]. In another case, the suspected mutation (A301S in the *rdl* gene) did not confer measurable resistance when introduced in *Plutella xylostella*, suggesting that insecticide resistance is not due to this mutation alone [20]. In a tour de force association study in *Bombyx mori*, four genes were found to contribute to coloration, egg diapause and two silk traits, respectively, and the phenotypic effect of three of these genes was confirmed by CRISPR/Cas9 [21].

The rapid explosion of whole genome sequencing combined with CRISPR/Cas9 technology is now broadening the range of insect species and phenotypic traits that can be explored to identify evolutionary-relevant mutations.

### Cis-regulatory and coding mutations are partitioned differently

Previous publications highlighted that Gephebase contains a higher number of coding mutations than cis-regulatory changes for all Eukaryotes [7,8,22]. We find here that this trend is also true for insects (Fig. 2C). Noticeably, only 3% of the 412 insect coding mutations present in Gephebase are curated as “Unknown” because the exact causal nucleotide change has not been pinpointed. In contrast, 59% of the 180 insect cis-regulatory mutations are curated as “Unknown”. This is certainly related to the fact that coding changes are easier to study than cis-regulatory mutations. As a result, our genetic knowledge is biased towards coding changes [7,8,22].

When all insects and traits are considered together, the rate of discovery of coding and cis-regulatory mutations does not appear to change over the years (Fig. 2C). As noted previously for all Eukaryotes [8], cis-regulatory mutations are more often found for morphological variation than for physiological variation in insects (Fig. 3A). This is especially clear when Coloration, implicating mostly cis-regulatory changes (89/108), is compared to Xenobiotic resistance, involving 352 coding mutations and 35 cis-regulatory mutations. This trend is also observed when removing potential bias due to the candidate gene approach, i.e. when considering only mutations identified by linkage and association mapping (Fig. 3A). Xenobiotic resistance arises through two main mechanisms, insensitivity of the target protein and enhanced metabolic detoxification [23,24]. In general, the former involves coding mutations at the target site and the latter involves gene amplification or cis-regulatory mutations at the detoxifying enzyme gene locus. Thus, genes encoding insecticide targets, such as *ace, para*, or *rdl*, are more likely to exhibit coding changes whereas genes encoding detoxifying enzymes such as P450, GST or esterase tend to show gene amplification or cis-regulatory changes (Fig. 3B).

**Fig. 3.**
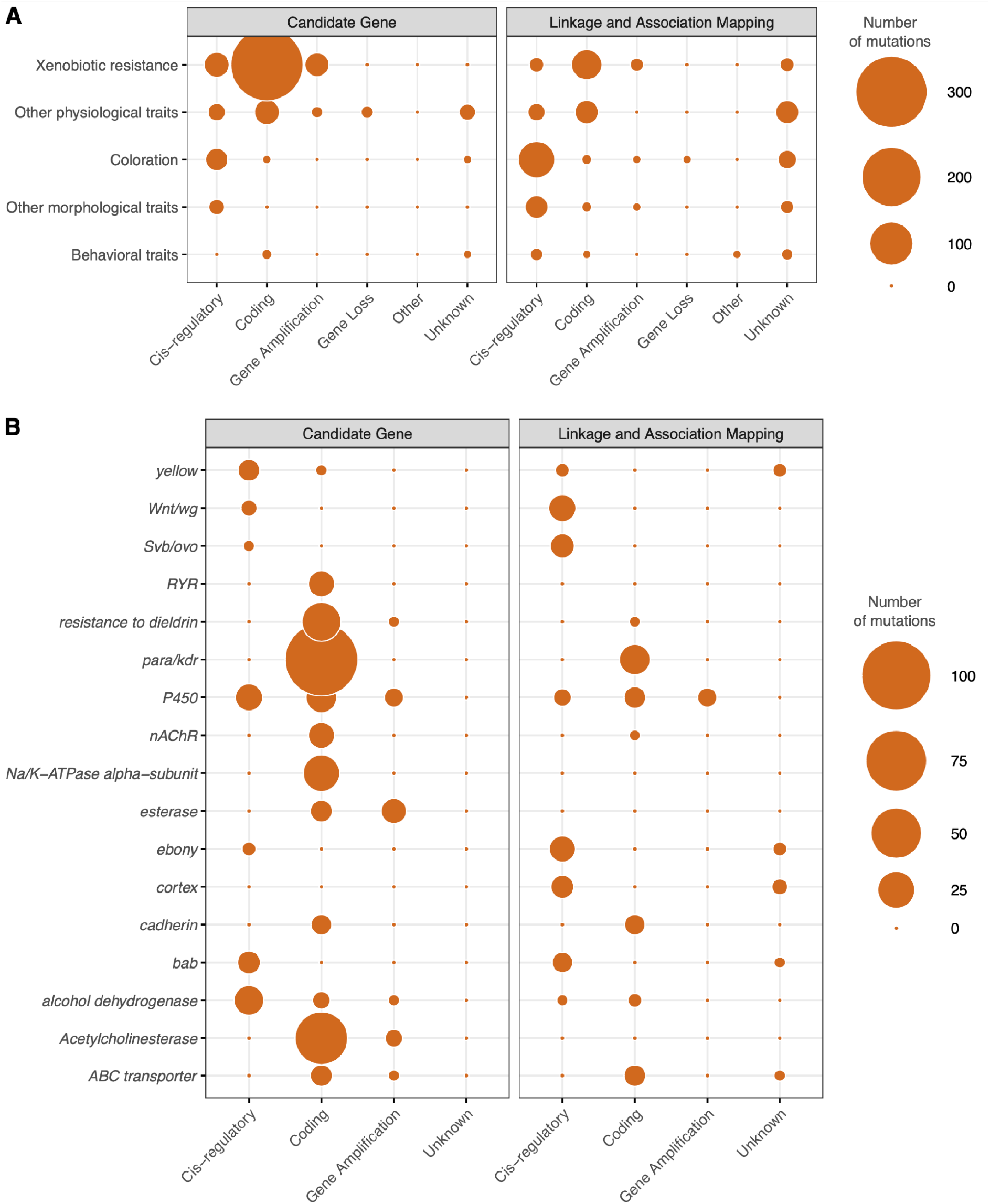
Number of mutations present in Gephebase according to molecular type, phenotypic trait, hotspot gene family, and experimental evidence. Mutations identified by a candidate gene approach are shown on the left and those found by association and linkage mapping on the right. Circle areas are proportional to the number of mutations present in Gephebase for insects in each category. ‘Other’ represents chimeric genes and supergenes. ‘Unknown’ indicates that the exact mutation was not identified. (A) Distribution of all the mutations reported in insects in Gephebase. (B) Distribution of the mutations for the top hotspot gene families (where more than 10 independent mutations were reported for each gene family).

Overall, the rate of discovery of coding versus cis-regulatory changes appears to depend on the trait of interest, the experimental approaches employed, and the gene loci identified.

### A trove of hotspot genes in insects

Many of the same insecticides and the same crops are used around the world, with intense global trade routes and long distance transport, so it is not surprising that the same new adaptive traits have arisen in many insects all over the world. What has been more surprising is that the mutations underlying natural adaptation have occurred multiple times independently in the same genes (so-called “hotspot genes”) in diverse populations and species [25]. Indeed, Gephebase reveals many hotspot genes in insects. The record is for the voltage sensitive sodium channel gene *vssc*/*para*, with more than 100 independent mutations that contribute to pyrethroid resistance across more than 40 insect species (File S1, Fig. 3A) [10]. In the aphid *Myzus persicae*, the ATG codon encoding for methionine at amino acid position 918 (sensitive allele) has undergone at least five independent changes (>ATT, Ile, >ATA, Ile, >CTG, Leu, >TTG, Leu and >ACG,Thr), all leading to pyrethroid resistance [26]. In the cotton bollworm, *Helicoverpa armigera*, at least eight distinct alleles of *CYP337B3* confer insecticide resistance and they seem to have arisen independently in diverse geographic locations [27]. One of them also introgressed into the native sibling species *Helicoverpa zea* [27]. Recent studies suggest that many mutations in hotspot genes that have increased in frequency due to insecticide selection over the last 50 years did not appear *de novo* but instead were already present at low frequency as standing genetic variation when insecticides were first introduced [26–28].

In other instances, mutations accumulated sequentially at the same gene locus, with each mutation contributing to incremental phenotypic change [29]. The *Cyp6g1* gene illustrates such a haplotype constructed step by step, through gene duplication and successive insertions of transposable elements, with intermediate haplotypes still found in natural populations [30]. Interestingly, it is in insects that most cases of such intra lineage hotspot genes are known. To our knowledge, in non-insect eukaryotes, only six genes have been reported to harbor 4 or more mutations, each contributing to variation in the phenotype of interest (see Gephebase for references). In contrast, 13 such cases are described for insects in Gephebase: *svb/ovo, alcohol dehydrogenase* in *D. melanogaster, D. erecta* and *D. virilis, Cyp6g1* in *D. melanogaster, CYP6P4/9* in *A. funestus, ebony* in *D. melanogaster, ace-2* in *D. melanogaster, nvd* in *D. pachea, Na/K-ATPase alpha-subunit* in the beetle *Chrysochus auratus* and the butterfly *Danaus plexippus*. Furthermore, the top intralineage hotspot gene is *svb/ovo*, with 9 characterized cis-regulatory mutations that affect the pattern of trichomes on the surface of *Drosophila* larvae [31]. Taxa outside insects are far behind, with a maximum of 5 evolutionary-relevant mutations clustered within individual genes, in *hemoglobin, HLA*, and *agouti* (see Gephebase for references).

Insect studies also highlight the important role of recombination in reshuffling alleles and creating new combinations conferring increased fitness. For example, the dark *ebony* allele in Uganda populations of *D. melanogaster* arose from the assembly of several mutations present as standing variation followed by two recent substitutions [32]. In the mosquito *A. aegypti*, analysis of sequences adjacent to the causal mutations showed that a 410L+1534C haplotype that confers pyrethroid resistance resulted from recombination between distinct 410L and 1534C alleles [33]. Conversely, inversions, which prevent recombination, are a common mechanism for maintaining large haplotypes that cause phenotypic divergence [34–37].

Several phenomena may explain why clusters of evolutionary-relevant mutations in single genes tend to be observed more frequently in insects than in other taxonomic groups. First, genetic studies may be easier in smaller organisms such as insects (compared to mice and zebrafish) and may facilitate the discovery of clustered mutations. Second, compared to animals and plants, insects generally have larger population sizes, allowing for more mutations to be present as standing variation, and shorter generation time, which may lead to faster response of populations to selection [28,38].

Finally, it is also in insects that the most complex polygenic architectures have been deciphered at the gene level. For example, the difference in abdominal pigmentation between the sister species *Drosophila santomea* and *Drosophila yakuba* is due to cis-regulatory mutations in five genes, *Abdominal-B, pdm3, tan, yellow* and *bab*, which interact in a well-characterized gene network [39].

In summary, recent genetic studies of insect phenotypic evolution highlight the important role of standing variation, hotspot genes, recombination, and introgression in the emergence of new traits in populations.

## Conclusions

Insects have been leading models in evolutionary genetics research for more than 100 years. Whereas previous studies relied on a handful of model systems, scientists are now starting to collect observations from many non-model systems. These studies will undoubtedly broaden our understanding of evolutionary processes and may help to develop novel approaches for sustainable agriculture, ecology, and public health.

## Abbreviations

CRISPR: Clustered Regularly Interspaced Short Palindromic Repeats
GST: Glutathione S-Transferase
QTL: Quantitative Trait Locus

## Supplementary materials

File S1. Table of all the insect entries extracted from Gephebase in November 2022 that were used for the present review.

(https://www.gephebase.org/search-criteria?/and+Taxon%20and%20Synonyms=Insecta/and+All=/and+splitMutations=true)

File S2. R script used to prepare all the figures.

## Acknowledgements

I am grateful to the anonymous and non-anonymous colleagues who suggested new studies to be curated into Gephebase via the Gephebase online tool. I greatly thank Séverine Wiltgen for Gephebase maintenance. I also thank the Ecole Normale Supérieure 2022 Master 1 students for discussions and oral presentations regarding the insect data present in Gephebase, and Arnaud Martin and David L. Stern for comments on a previous draft.

*at least 10% of your references should be selected and annotated as being papers of special interest (•) or outstanding interest (••). Annotated references MUST be from the past two years, and the annotation should provide a brief description of the major findings and the importance of the study*.

## References

1. Sturtevant AH: Genetic studies on Drosophila simulans. I. Introduction. Hybrids with Drosophila melanogaster. Genetics 1920, 5:488.

2. Rubin GM: Drosophila melanogaster as an experimental organism. Science 1988, 240:1453–1459.

3. Telis N, Lehmann BV, Feldman MW, Pritchard JK: A Bibliometric History of the Journal GENETICS. Genetics 2016, 204:1337–1342.

4. Li F, Zhao X, Li M, He K, Huang C, Zhou Y, Li Z, Walters JR: Insect genomes: progress and challenges. Insect Mol Biol 2019, 28:739–758.

5. Poelchau M, Childers C, Moore G, Tsavatapalli V, Evans J, Lee C-Y, Lin H, Lin J-W, Hackett K: The i5k Workspace@ NAL—enabling genomic data access, visualization and curation of arthropod genomes. Nucleic Acids Res 2015, 43:D714–D719.

6. Mei Y, Jing D, Tang S, Chen X, Chen H, Duanmu H, Cong Y, Chen M, Ye X, Zhou H: InsectBase 2.0: a comprehensive gene resource for insects. Nucleic Acids Res 2022, 50:D1040–D1045. Paper describing the database InsectBase 2.0 (http://v2.insect-genome.com/), which covers 815 insect genomes and provides JBrowse2, Synteny Viewer and BLAST services.

7. Courtier-Orgogozo V, Arnoult L, Prigent SR, Wiltgen S, Martin A: Gephebase, a database of genotype–phenotype relationships for natural and domesticated variation in Eukaryotes. Nucleic Acids Res 2020, 48:D696–D703. Description of the database Gephebase, which compiles published data about the genes and the mutations contributing to natural evolution and domestication in Eukaryotes.

8. Stern DL, Orgogozo V: The loci of evolution: How predictable is genetic evolution ? Evol Int J Org Evol 2008, 62:2155–2177.

9. Feyereisen R, Dermauw W, Van Leeuwen T: Genotype to phenotype, the molecular and physiological dimensions of resistance in arthropods. Pestic Biochem Physiol 2015, 121:61–77.

10. Dong K, Du Y, Rinkevich F, Nomura Y, Xu P, Wang L, Silver K, Zhorov BS: Molecular biology of insect sodium channels and pyrethroid resistance. Insect Biochem Mol Biol 2014, 50:1–17.

11. Nauen R, Bass C, Feyereisen R, Vontas J: The role of cytochrome P450s in insect toxicology and resistance. Annu Rev Entomol 2022, 67:105–124. Recent review of the role of insect cytochrome P450 monooxygenases in the detoxification of xenobiotics.

12. Jurat-Fuentes JL, Heckel DG, Ferré J: Mechanisms of resistance to insecticidal proteins from Bacillus thuringiensis. Annu Rev Entomol 2021, 66:121–140. Recent review of the mechanisms of resistance to Bt toxins, with emphasis on field-evolved resistance and key resistance genes.

13. McCulloch KJ, Macias-Muñoz A, Briscoe AD: Insect opsins and evo-devo: what have we learned in 25 years? Philos Trans R Soc B 2022, 377:20210288.

14. Massey JH, Wittkopp PJ: The genetic basis of pigmentation differences within and between Drosophila species. Curr Top Dev Biol 2016, 119:27–61.

15. Elkin J, Martin A, Courtier-Orgogozo V, Santos ME: Meta-analysis of the genetic loci of pigment pattern evolution in vertebrates. bioRxiv 2022,

16. Stork NE: How many species of insects and other terrestrial arthropods are there on Earth. Annu Rev Entomol 2018, 63:31–45.

17. Gantz VM, Akbari OS: Gene editing technologies and applications for insects. Curr Opin Insect Sci 2018, 28:66–72.

18. Douris V, Denecke S, Van Leeuwen T, Bass C, Nauen R, Vontas J: Using CRISPR/Cas9 genome modification to understand the genetic basis of insecticide resistance: Drosophila and beyond. Pestic Biochem Physiol 2020, 167:104595. •• Thoughtful review of the advantages and limitations of using CRISPR/Cas9 genome modifications to explore the genetic basis of insecticide resistance, illustrated by several recent studies.

19. Jin L, Wang J, Guan F, Zhang J, Yu S, Liu S, Xue Y, Li L, Wu S, Wang X, et al.: Dominant point mutation in a tetraspanin gene associated with field-evolved resistance of cotton bollworm to transgenic Bt cotton. Proc Natl Acad Sci U S A 2018, 115:11760–11765.

20. Guest M, Goodchild JA, Bristow JA, Flemming AJ: RDL A301S alone does not confer high levels of resistance to cyclodiene organochlorine or phenyl pyrazole insecticides in Plutella xylostella. Pestic Biochem Physiol 2019, 158:32–39.

21. Tong X, Han M-J, Lu K, Tai S, Liang S, Liu Y, Hu H, Shen J, Long A, Zhan C, et al.: High-resolution silkworm pan-genome provides genetic insights into artificial selection and ecological adaptation. Nat Commun 2022, 13:5619. •• Tour de force study of more than 1000 new high-resolution genome sequences of the silkworm Bombyx mori, with identification of four genes underlying classical silkworm mutants and their validation by CRISPR/Cas-9.

22. Stern D, Orgogozo V: Is Genetic Evolution Predictable? Science 2009, 323:746–751.

23. Bass C, Field LM: Gene amplification and insecticide resistance. Pest Manag Sci 2011, 67:886–890.

24. Bass C, Puinean AM, Zimmer CT, Denholm I, Field LM, Foster SP, Gutbrod O, Nauen R, Slater R, Williamson MS: The evolution of insecticide resistance in the peach potato aphid, Myzus persicae. Insect Biochem Mol Biol 2014, 51:41–51.

25. Martin A, Orgogozo V: The Loci of repeated evolution: a catalog of genetic hotspots of phenotypic variation. Evol Int J Org Evol 2013, 67:1235–1250.

26. Singh KS, Cordeiro EMG, Troczka BJ, Pym A, Mackisack J, Mathers TC, Duarte A, Legeai F, Robin S, Bielza P, et al.: Global patterns in genomic diversity underpinning the evolution of insecticide resistance in the aphid crop pest Myzus persicae. Commun Biol 2021, 4:847. •• Beautiful study of the emergence and spread of several insecticide resistance in the aphid crop pest Myzus persicae, involving pre-adaptation to plant toxic compounds and repeated evolution at specific genomic loci.

27. Walsh TK, Joussen N, Tian K, McGaughran A, Anderson CJ, Qiu X, Ahn S-J, Bird L, Pavlidi N, Vontas J, et al.: Multiple recombination events between two cytochrome P450 loci contribute to global pyrethroid resistance in Helicoverpa armigera. PloS One 2018, 13:e0197760.

28. Hawkins NJ, Bass C, Dixon A, Neve P: The evolutionary origins of pesticide resistance. Biol Rev 2019, 94:135–155.

29. Scott JG: Life and death at the voltage-sensitive sodium channel: evolution in response to insecticide use. Annu Rev Entomol 2019, 64:243–257.

30. Schmidt JM, Good RT, Appleton B, Sherrard J, Raymant GC, Bogwitz MR, Martin J, Daborn PJ, Goddard ME, Batterham P: Copy number variation and transposable elements feature in recent, ongoing adaptation at the Cyp6g1 locus. PLoS Genet 2010, 6:e1000998.

31. Stern DL, Frankel N: The structure and evolution of cis-regulatory regions: the shavenbaby story. Philos Trans R Soc Lond B Biol Sci 2013, 368:20130028.

32. Rebeiz M, Pool JE, Kassner VA, Aquadro CF, Carroll SB: Stepwise modification of a modular enhancer underlies adaptation in a Drosophila population. Science 2009, 326:1663–1667.

33. Fan Y, O’Grady P, Yoshimizu M, Ponlawat A, Kaufman PE, Scott JG: Evidence for both sequential mutations and recombination in the evolution of kdr alleles in Aedes aegypti. PLoS Negl Trop Dis 2020, 14:e0008154.

34. Ando T, Niimi T: Development and evolution of color patterns in ladybird beetles: A case study in Harmonia axyridis. Dev Growth Differ 2019, 61:73–84.

35. Gutiérrez-Valencia J, Hughes PW, Berdan EL, Slotte T: The genomic architecture and evolutionary fates of supergenes. Genome Biol Evol 2021, 13:evab057.

36. Fuller ZL, Koury SA, Phadnis N, Schaeffer SW: How chromosomal rearrangements shape adaptation and speciation: Case studies in Drosophila pseudoobscura and its sibling species Drosophila persimilis. Mol Ecol 2019, 28:1283–1301.

37. Wang S, Teng D, Li X, Yang P, Da W, Zhang Y, Zhang Y, Liu G, Zhang X, Wan W: The evolution and diversification of oakleaf butterflies. Cell 2022, 185:3138–3152. •• Successful association mapping study in the oakleaf butterfly, an iconic non-model species, where the different wing morphs are shown to be determined by the wing patterning gene cortex, which has been maintained in the populations by long-term balancing selection.

38. Karasov T, Messer PW, Petrov DA: Evidence that adaptation in Drosophila is not limited by mutation at single sites. PLoS Genet 2010, 6:e1000924.

39. Liu Y, Ramos-Womack M, Han C, Reilly P, Brackett KL, Rogers W, Williams TM, Andolfatto P, Stern DL, Rebeiz M: Changes throughout a Genetic Network Mask the Contribution of Hox Gene Evolution. Curr Biol 2019,

